# Prolonged podocyte depletion in larval zebrafish resembles mammalian focal and segmental glomerulosclerosis

**DOI:** 10.1101/706267

**Authors:** Kerrin Ursula Ingeborg Hansen, Florian Siegerist, Sophie Daniel, Maximilian Schindler, Antje Blumenthal, Weibin Zhou, Karlhans Endlich, Nicole Endlich

## Abstract

Although FSGS has been in the scientific focus for many years, it is still a massive burden for patients with no causal therapeutic option. In FSGS, podocytes are injured, parietal epithelial cells (PECs) are activated and engage in the formation of cellular lesions leading to progressive glomerular scarring. Herein we show that podocyte-depleted zebrafish larvae develop acute proteinuria, severe foot process effacement and activate PECs which create cellular lesions and deposit extracellular matrix on the glomerular tuft. We therefore propose that this model shows features of human FSGS and show its applicability for a high-throughput drug screening assay.

## Introduction

Podocyte-depletion has been shown one of the initial steps in the development of glomerulosclerosis.^1^ In focal and segmental glomerulosclerosis (FSGS), podocytes are injured and parietal epithelial cells (PECs) are activated thus forming cellular lesions on the glomerular tuft.^2^ Recently, Kuppe et al. showed that upon activation, PECs develop a cuboidal phenotype and are more prone to contribute to sclerotic lesions.^3^ Since experimental procedures in rodents are not ideally suitable for high-throughput drug screening assays, we and others used the larval zebrafish pronephros as a vertebrate model^4^ to study glomerular morphology^5^ and permselectivity of the glomerular filtration barrier.^6^ At 48 hours past fertilization, zebrafish develop a single glomerulus attached to a pair of tubules.^4^ As in mammals, the glomerular filtration barrier consists of a fenestrated endothelium, the glomerular basement membrane (GBM) and interdigitating podocytes bridged by a conserved slit diaphragm.^4^ As an injury model we used the nitroreductase/metronidazole (NTR/MTZ) model for podocyte depletion, since the prodrug MTZ is activated exclusively in podocytes expressing the NTR under control of the *nphs2* promotor leading to rapid onset of proteinuria and edema resembling human nephrotic syndrome.,^7,5,6^ The aim of this study was to examine the glomerular response upon mild podocyte depletion and to investigate the applicability of this model for human FSGS.

## Results

To generate conditions of prolonged podocyte depletion, we treated *Cherry* larvae (Tg(*nphs2*:GAL4); Tg(UAS:Eco.nfsb-mCherry); mitfa ^w2/w2^) with either low-dose MTZ (80 µmol L^-1^) or 0.1% DMSO for 48 hours, beginning at 4 days past fertilization (dpf). Edema was graded into four categories (Fig.1A). MTZ-treated larvae developed edema one day after washout. Until 8 dpf, 69% (of n=275) of living MTZ-treated larvae showed medium or severe edema formation (Fig. 1B). 51% of MTZ-treated larvae died at 9 dpf (Fig.1C). In contrast, 10% (of n=251) of control-treated larvae showed edema of any severity or died during the considered period of time (Fig.1B, C).

**Fig. 1.**
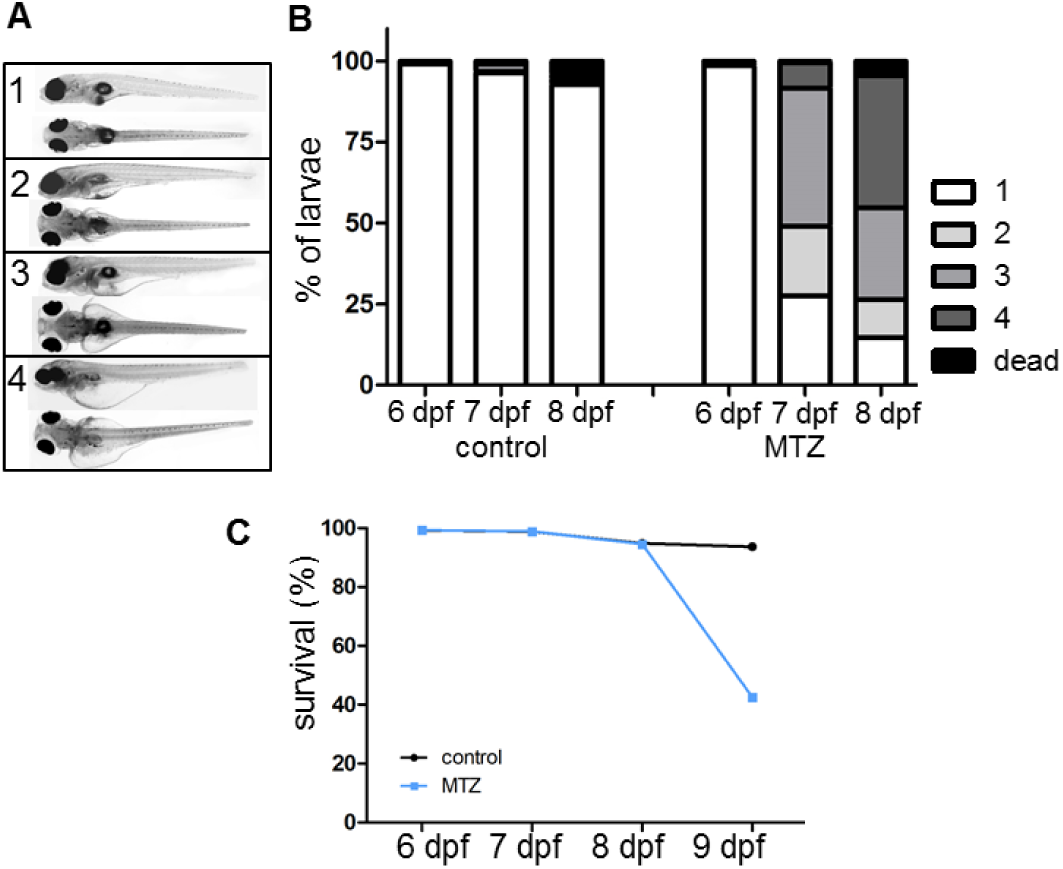
Grading of edema and survival following podocyte depletion. Figure 1A shows representative micrographs of 8 dpf zebrafish larvae to demonstrate grading of developing edema after podocyte depletion. (1) no edema, (2) mild edema, (3) medium edema, (4) severe edema. The graph in B shows relative distributions of phenotypes in n=251 control-treated and n=275 MTZ-treated larvae in four individual experiments. As visualised in the diagram, survival rates of podocyte-depleted larvae are similar to controls until an abrupt decrease at 9 dpf.

To quantify proteinuria, we intravenously injected 10 kDa (TRITC) and 500 kDa (FITC) dextran directly after MTZ-treatment and measured the intravascular fluorescence after 19 hours (Fig.2A). The analysis of FITC-fluorescence in the caudal vein showed a decrease of 59% in MTZ-treated larvae after 19 hours (p=0.0002 compared to controls; n=31); whereas fluorescence in controls did only slightly decline during the considered period (Fig. 2B, Suppl.Fig.1).

**Fig. 2.**
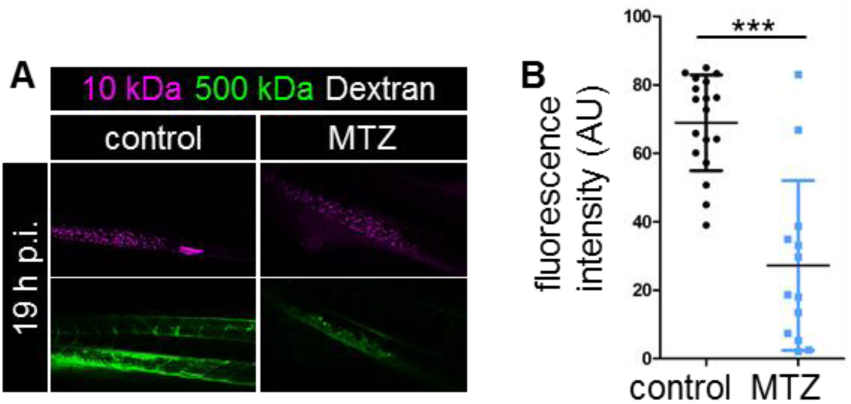
Functional analysis of glomerular filtration barrier. Panel A shows confocal laser scanning micrographs of podocyte-depleted and control larvae 19h after injection with a mixture of FITC-conjugated 500kDa and Alexa 647-conjugated 10kDa dextran directly after metronidazole washout. Notice the decrease of 500kDa dextran in podocyte-depleted larvae, whereas 10kDa dextran accumulates in the tissue. As shown in graph B, fluorescence intensity of FITC measured in the caudal vein was significantly lower in podocyte-depleted larvae 19 h after dextran injection (see Suppl.Fig. 1 for full data).

Transmission electron microscopy (TEM) at 9 dpf showed foot process effacement of remaining podocytes in MTZ-treated larvae, whereas controls exhibited regularly shaped foot processes (Fig.3A). Immunostaining showed a reduction of *podocin* in MTZ-treated larvae in comparison to controls, which displayed a regular staining pattern along the glomerular capillaries (Fig.3B).

**Fig. 3.**
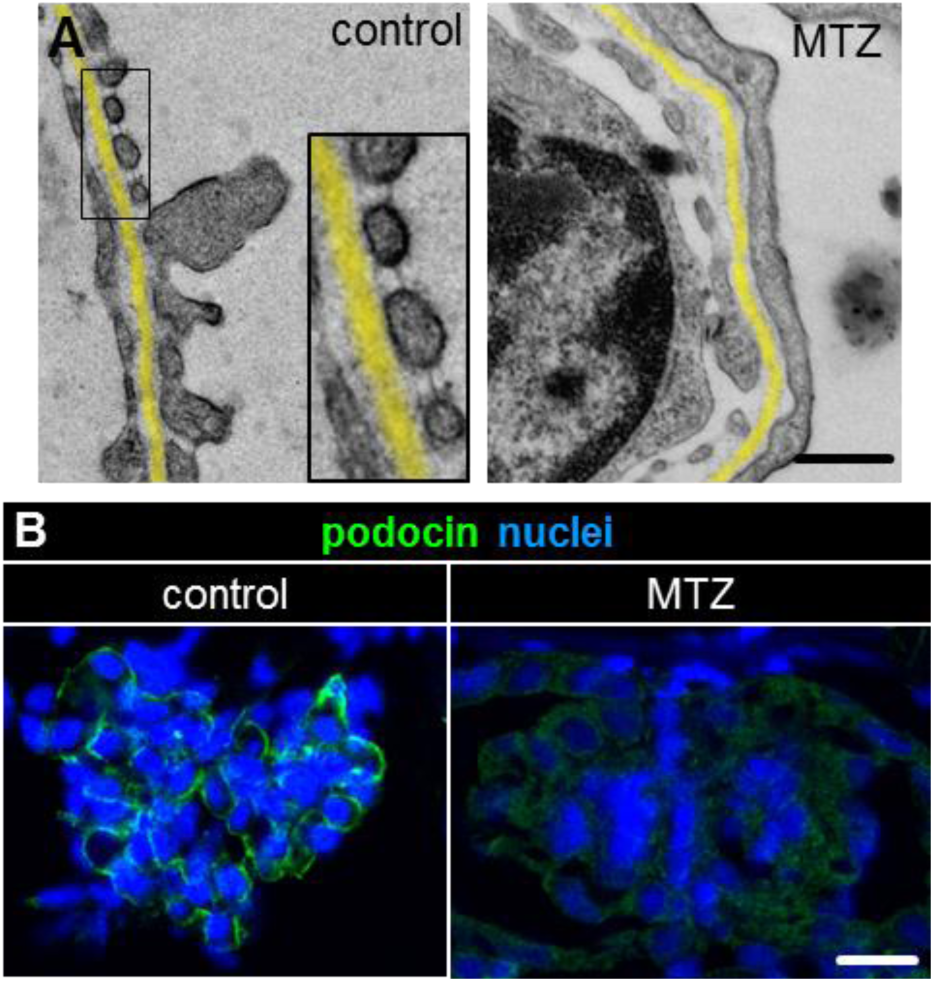
Structural analysis of the glomerular filtration barrier. Transmission electron micrographs of 9 dpf control larvae in panel A reveal a normal morphology with fine foot processes connected by a slit diaphragm whereas podocyte foot processes in MTZ-treated larvae are broadly effaced. The GBM is highlighted in yellow. Scale bar represents 400 nm. Panel B shows the linear staining pattern of *podocin* three days after vehicle treatment in controls while the signal is greatly reduced in podocyte-depleted larvae. The scale bar represents 25 µm.

Histological analysis showed a reduced glomerular cell density in MTZ-treated larvae compared to controls (median: 0.014 nuclei per µm^2^ in MTZ-treated larvae versus 0.022 nuclei per µm^2^; p=0.0011; n=24) (Suppl.Fig.2A, B, C). It remained reduced over three days of regeneration (0.0161 nuclei per µm^2^ versus 0.0325 nuclei per µm^2^ in controls at 9 dpf; p=0.0003; n=24), although absolute cell numbers at the capillary tuft of MTZ-treated larvae increased to a level similar to controls (Suppl.Fig.2D). Numbers of podocytes per glomerular cross-section determined by TEM remained significantly reduced (mean: 9.0 versus 13.25 at 9 dpf; p=0.0043; n=17) (Suppl.Fig.2E). Bowman’s capsules were enlarged in MTZ-treated larvae determined as the area of the largest glomerular cross-section for every larva. At 6 dpf, the mean was 1043 µm^2^ after MTZ-treatment compared to 538 µm^2^ in controls (Suppl.Fig.2C, p=0.0008, n=24). The difference was even higher at 9 dpf (1085 µm^2^ in MTZ-treated larvae versus 374 µm^2^, p=0.0019, n=24). In podocyte-depleted larvae, distinct severities of podocyte impairment could be discriminated by TEM: In 45% (of n=11) of MTZ-treated larvae, glomeruli showed a GBM with a uniform electron density and as well fenestrated endothelial cells similar to controls, but with severe foot process effacement (Fig.4A). Only few capillaries showed intact foot processes. Frequently, dilatations of the subpodocyte space were found in podocyte-depleted larvae (asterisk in Fig.4A). In 36% of MTZ-treated larvae, no typical foot processes of podocytes were visible. As shown in Fig.4B, visceral epithelial cells were instead showing tight junctions and microvillous transformation. Total numbers of local electron-dense contacts to neighboring cells per visceral epithelial cell were increased from 0.29 in controls to 2.2 tight junctions per cell in MTZ-treated larvae (p=0.0002, n=20) (Fig.4B, C).

**Fig. 4.**
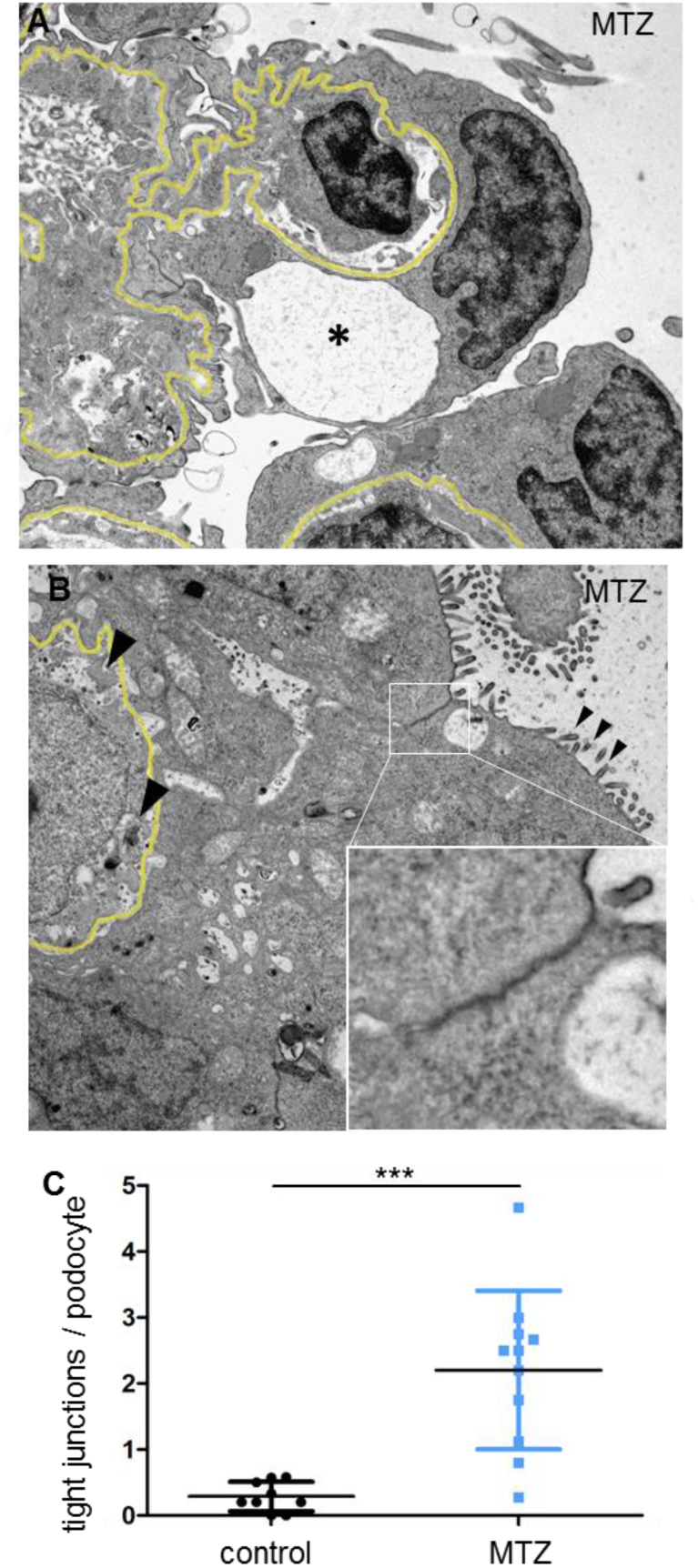
Ultructural analysis of podocytes and PECs. As shown in the representative micrograph in A, podocyte-depleted glomeruli displayed signs of podocyte impairment: Adjacent to a capillary displaying almost normal morphology; podocyte foot process effacement and subpodocyte space pseudocys (asterisk) can be seen. MTZ-treated larvae showed markedly increased tight junctions between remaining podocytes in n=20 larvae (B, C). An electron-dense tight junction is shown in detail in the bottom right corner of the picture. Black arrowheads accentuate loss of fenestration and cellular integrity of the capillary endothelium. Scale bar represents 2 µm.

Histomorphologic analysis showed a thickening of the PEC-layer that progressed between 6 and 9 dpf (Fig.5A). After MTZ washout (6 dpf), mean height of PECs was 0.9 µm in controls and 1.11 µm in podocyte-depleted larvae (p=0.0043; n=24). At 9 dpf, it was 0.78 µm in controls and 3.96 µm after MTZ treatment (p<0.0001; n=24) (Fig.5B).

**Fig. 5.**
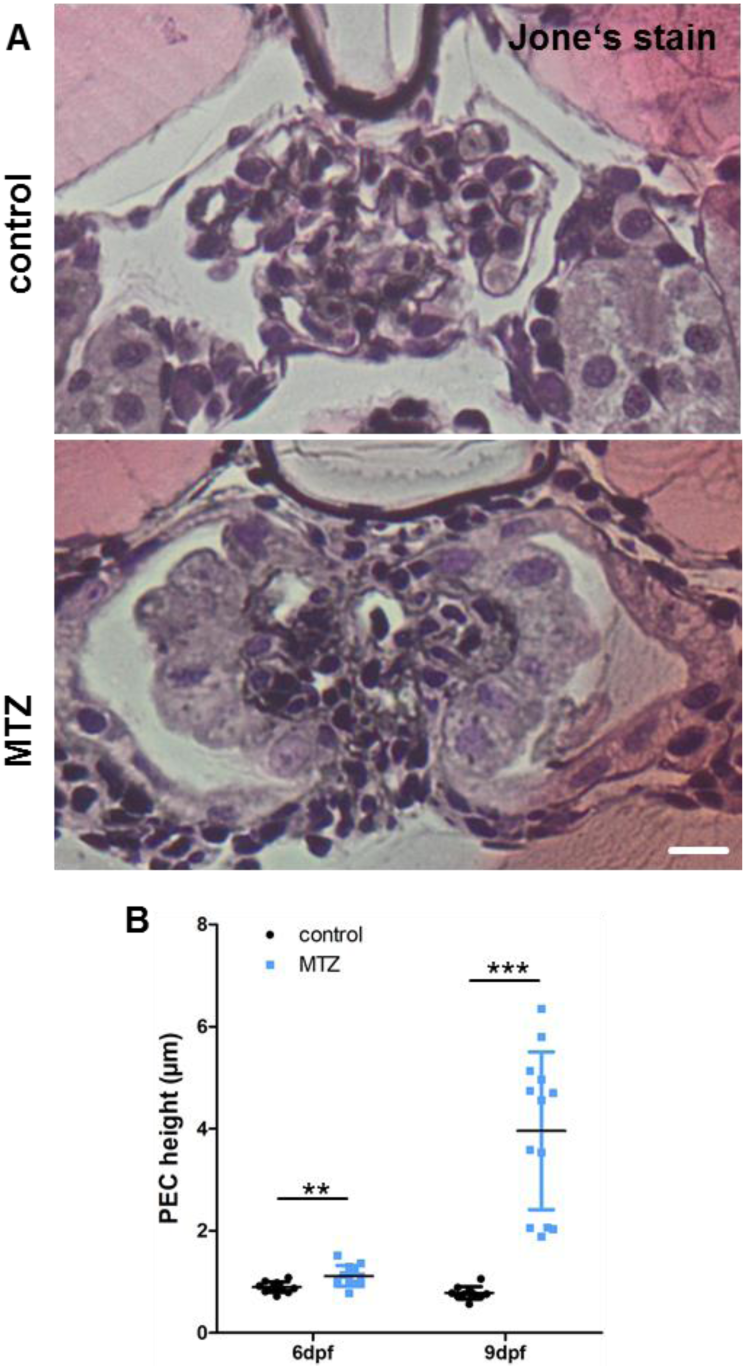
Morphometry of PECs following podocyte depletion. Panel A shows representative micrographs of Jones-stained plastic sections of podocyte-depleted and control larvae. The scale bar represents 10 µm. The graph in B shows the result of PEC height measurements in H&E-stained plastic sections of n=25 podocyte-depleted and n=23 control larvae. A statistically significant increase is visible in podocyte-depleted larvae directly after, and three days after treatment.

Immunostaining for *proliferating cell nuclear antigen* (*pcna*), revealed proliferating PECs only in podocyte-depleted larvae (Fig.6A).

**Fig. 6.**
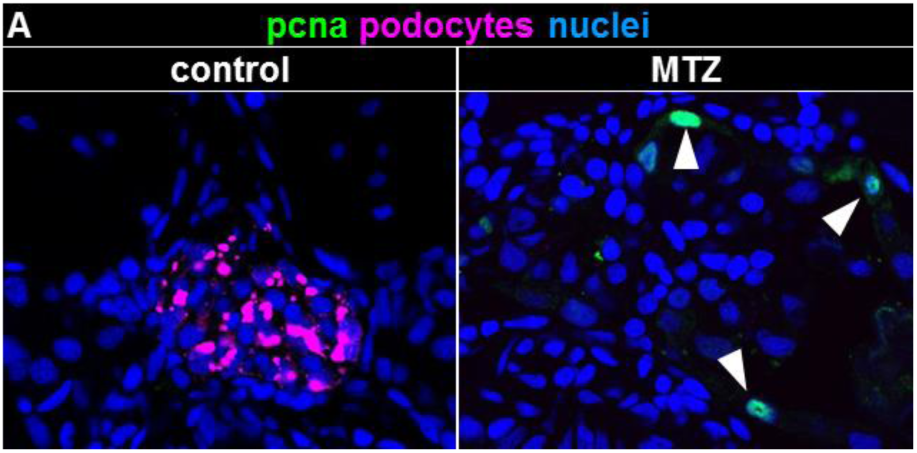
Analysis of PEC proliferation. The micrographs in panel C show *pcna*-staining of larvae collected at 9 dpf. Podocyte-depleted larvae show *pcna*-positivity of PECs (white arrowheads). Scale bar represents 10 µm.

Furthermore, podocyte-depleted larvae frequently showed adhesions between the parietal and the visceral glomerular cell layer visible in H&E sections (Fig.7A) and TEM micrographs (Fig.7B). Since podocyte loss in humans is followed by progressive scarring of the glomerulus, we investigated extracellular matrix deposition in podocyte-depleted zebrafish larvae. While Jone’s stain showed minimal accumulation of silver-positive material on the glomerular tuft at 9 dpf (Fig.5A), a strong accumulation of laminin was seen on the glomerular tuft (Fig.2F, Suppl.Fig.4) with significant GBM-thickening (Fig.8A,C). However, we could not detect accumulation of collagen I at 9 dpf (Suppl.Fig.3). To validate that the cuboidal cells on the glomerular tuft were of PEC-origin, we immunostained for *pax2a* which is a marker of the tubular neck segment.^8^ PECs where strongly *pax2a*-positive under baseline conditions as well as the cuboidal cells on the glomerular tuft (Fig.8B, Suppl.Fig.5).

Using histology and TEM, we additionally found neutrophils and macrophages within Bowman’s capsule only in podocyte-depleted larvae (Fig. 2I and Suppl.Fig.6).

**Fig. 7.**
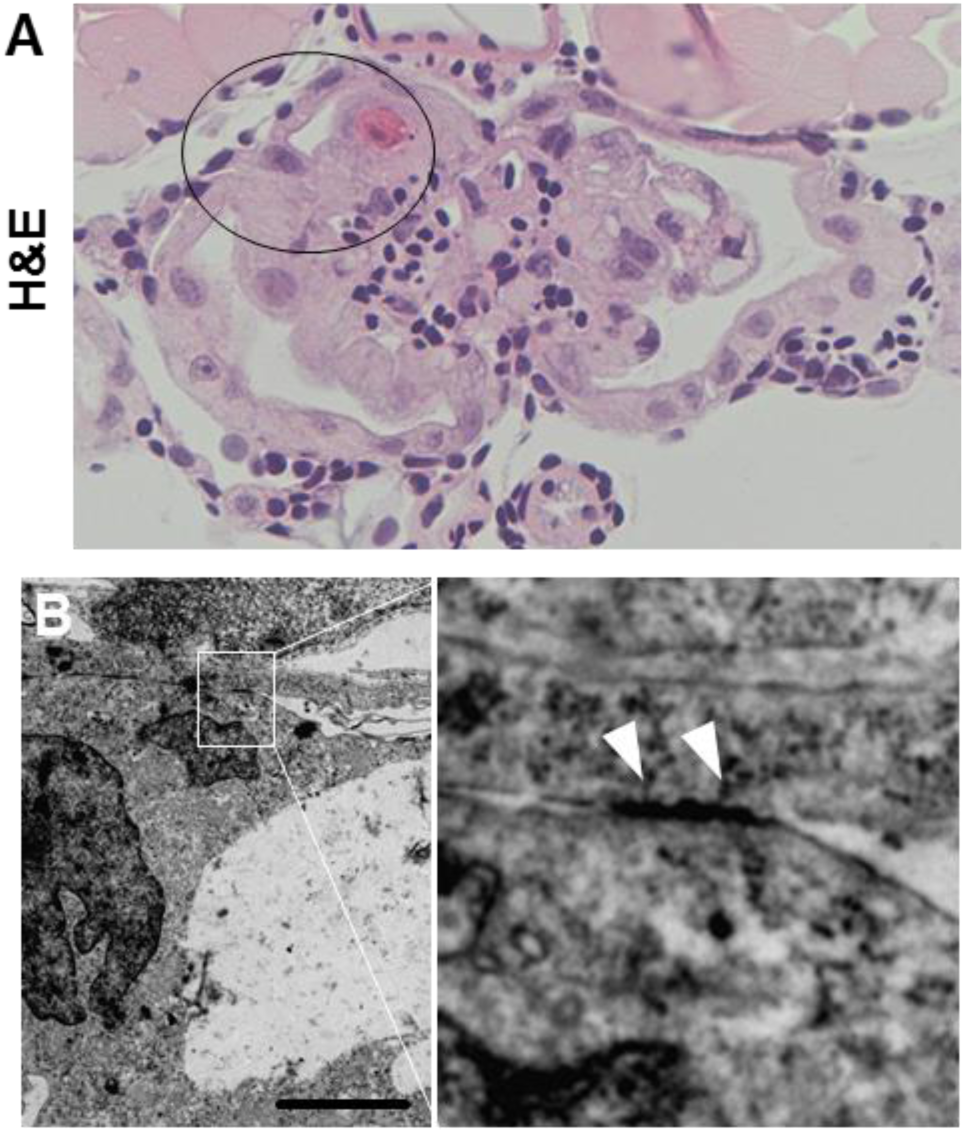
Structural analysis of parietovisceral adhesions. Picture A shows parieto-visceral adhesion in an H&E-stained plastic section. Scale bar represents 20µm. The transmission electron micrograph shown in picture B further characterises one adhesion. Scale bar represents 2 µm. White arrowheads highlight local electron-dense parieto-visceral contacts in the magnification of B.

**Fig. 8.**
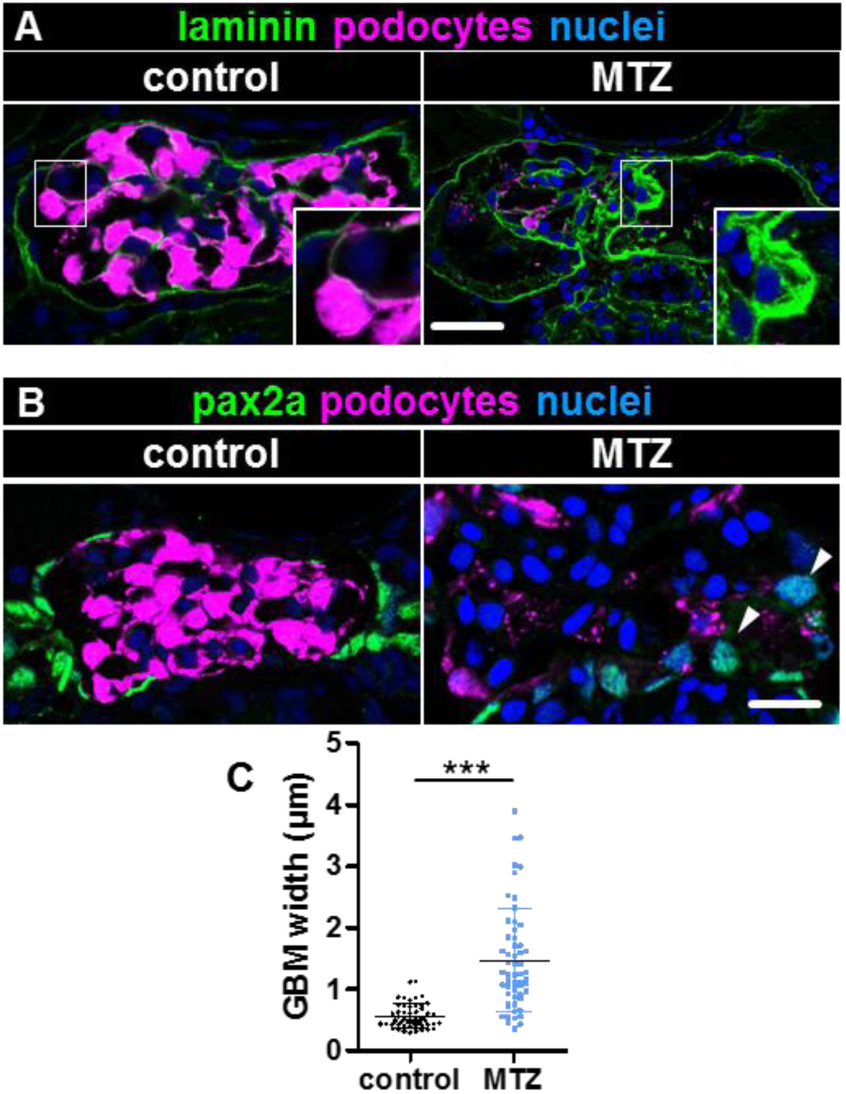
Novel glomerular cells accumulate extracellular matrix and are of PEC-origin. Panel A shows immunostaining for laminin in podocyte-depleted and control larvae at 9 dpf. A significant deposition of laminin on the glomerular tuft can be noticed as shown in the inserts. The graph in C quantifies the thickness of the laminin-layer within the GBM measured orthogonally as the full width at half maximum. Podocyte-depleted larvae showed statistically significant increased laminin-deposition as measured in 5 randomly-picked capillaries in each 3 consecutive glomerular cross-sections of n=5 podocyte-depleted and n=6 control larvae. Panel B demonstrates *pax2a*-positive PEC-nuclei under baseline conditions with increased expression of *pax2a* in cuboidal PECs and cuboidal cells on the glomerular tuft 3 days after podocyte depletion (arrowheads, overview in Suppl.Fig.5).

**Fig. 8.**
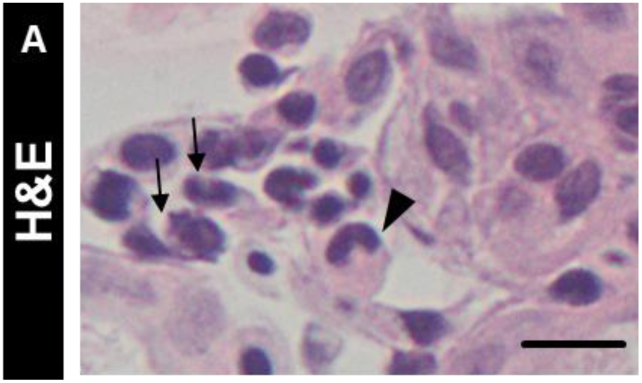
Phagocytic cells are recruited to the glomerulus following podocyte depletion. The black arrows in picture A mark neutrophils with the characteristic lobate nucleus. The black arrowhead marks the horseshoe-shaped nucleus of an intracapsular macrophage (overview in Suppl.Fig.6).

## Discussion

The small larval size and conserved morphology of the pronephros offers a general applicability for high-throughput assays. As shown before, the NTR/MTZ model is a valuable tool to investigate glomerular adaption and regeneration after podocyte loss by enabling specific podocyte-depletion.^7,5^ In this study, we adapted the injury so that larvae survived long enough to investigate glomerular response. As described before, MTZ-treated larvae developed periocular edema, which is a hallmark for proteinuria^7^ which we verified using clearance of high-molecular-weight dextran. After 3 days, remaining podocytes showed foot process effacement and reduction of *podocin* as a sign of podocyte injury. An increase of tight junctions between neighboring podocytes can be interpreted as a mechanism to prevent detachment from the GBM under increased mechanical forces and has been described in mammalian models.^9,10^ Although the size of the glomerular tuft increased, the number of podocytes remained significantly lower than in controls, indicating podocyte-hypertrophy. Previous work has shown that insufficient podocyte-hypertrophy and subsequent mechanical stress leads to the development of subpodocyte space pseudocysts and ultimately to podocyte detachment.^11^ Furthermore, increased filtration in the sub-podocyte space has been suggested to play an important role in regulation of glomerular permeability.^12^ Another key feature of mammalian FSGS is the activation of PECs which contribute to fibrotic lesions as it has been shown by Smeets and colleagues^13^ and thickening of Bowman’s capsule.^14^ In our model, PECs proliferated and changed their morphology towards a cuboidal, microvillous phenotype. Taken together, these findings support the hypothesis that upon podocyte-depletion, flat PECs transform to cuboidal PECs, as it has recently been shown by Kuppe et al.^3^ Together with the typical cytomorphology and the expression of *pax2a* and not *podocin*, we characterized them as cuboidal PECs that covered the denuded areas of the GBM in the most severely injured 36% of MTZ-treated larvae, which fits previous findings in a murine model of collapsing FSGS.^15^ In contrast to that, other groups have suggested, that PECs are recruited to the glomerular tuft and replace podocytes.^16,17^ Another work additionally mentions attachment of PECs to the apical sides of podocytes after activation and both propose extracellular matrix conglomerates in FSGS to be synthesized by PECs as we have demonstrated in our model.^15,18^ Although zebrafish larvae seem to have the general ability to develop sclerosis^19^, they seem to develop remarkably little fibrosis and instead regenerate tissue as shown for cardiomyocytes.^20^

Taken together, we show that podocyte-depletion in zebrafish resembles FSGS in important characteristics like proteinuria, development of edema, formation of viscero-parietal adhesions, PEC activation and proliferation as well as deposition of extracellular matrix. Our results establish a basis not only for the use of FSGS-like disease in zebrafish as a model for further studies investigating the pathogenesis of FSGS, but also for assessing the effects of potential drugs on disease development in a vertebrate model suitable for high-throughput experiments.

## Materials and Methodsxs

### Zebrafish husbandry and MTZ treatment

Zebrafish *(Danio rerio)* were bred, maintained and staged according as described before.^6^ We used the transparent and transgenic strain Tg(*nphs2*:GAL4); Tg(UAS:Eco.nfsb-mCherry); mitfa^w2/w2^ for all experiments. Larvae express the bacterial enzyme nitroreductase and the fluorescent protein *mCherry* exclusively in podocytes. All experiments were performed according to German animal protection law overseen by the “Landesamt für Landwirtschaft, Lebensmittelsicherheit und Fischerei, Rostock” of the federal state of Mecklenburg - Western Pomerania. For drug treatment, 0.1 % DMSO was freshly diluted in E3-embryo medium. MTZ (Sigma-Aldrich, St. Louis, MO, USA) was added at a concentration of 80 µmol L^-1^ for all experiments. Controls were treated with 0.1 % DMSO-solution only. Treatment was started at 4 dpf and a treatment period of 48 h was held for all experiments.

### Histology

Larvae for histological analysis were fixed at 6 dpf and at early 9 dpf in 4% paraformaldehyde at 4°C overnight. Plastic embedding was performed in Technovit^®^ 7100 (Kulzer GmbH, Hanau, Germany) as per manufacturer’s instructions. 4 µm-sections were made with a Jung RM 2055 rotational microtome (Leica Microsystems, Wetzlar, Germany). H&E and PAM silver staining according to Jones were performed adhering to Technovit^®^ 7100 routine staining protocols.

### Immunofluorescence staining

For *podocin* and *collagen I alpha 1* staining, larvae were fixed in 4% paraformaldehyde at 4°C overnight and embedded in paraffin according to standard protocols. 5 µm sections were made on a Leica SM 200R microtome. After heat-mediated antigen retrieval, sections were incubated with primary antibodies 1:500 rabbit anti-*podocin* (Proteintech, IL, USA) or 1:500 rabbit anti-*col1a1* (GeneTex, CA, USA) at 4°C overnight. For *pcna, pax2a* and *laminin* staining, larvae were fixed in 2% paraformaldehyde at 4°C overnight. 30% sucrose in PBS was mixed 1:1 with TissueTek (Sakura Finetek Europe, AV, Netherlands) and used for infiltration for 3 hours at room temperature. Samples were snap-frozen in liquid nitrogen. 5 µm sections were made on a Microm HM 560 microtome (Thermo Fisher Scientific, MA, USA). After permeabilization with 0.3% Triton X-100 and five washes with PBS, slides were incubated with 1:50 rabbit anti-*pcna* (sc-56, Santa Cruz Biotechnology, TX, USA), 1:500 rabbit anti-pax2a (ab229318, abcam) or 1:35 rabbit anti-*laminin* (L9393, Sigma-Aldrich) at 4°C overnight. Slides were washed five times in PBS. For all stainings, Alexa 488 or 647-conjugated goat anti-rabbit IgG F(ab)2 antibody fragment (Dianova, Hamburg, Germany) were used at 1:300 dilution. Nuclei were counterstained with 0.1 mg/ml DAPI (4′,6-Diamidine-2′-phenylindole dihydrochloride, Sigma, MO, USA) for 20 minutes. After one wash with PBS, slides were mounted with Mowiol for microscopy (Roth, Karlsruhe, Germany).

Immunofluorescence micrographs were acquired with a TCS SP5 confocal laser scanning microscope using the 63x, 1.4 NA oil immersion objective (Leica Microsystems, Wetzlar, Germany). Brightfield images were acquired with an Olympus BX50 light microscope using the 40x, 0.6 NA objective (Olympus, Hamburg, Germany). ImageJ V1.51f (Wayne Rasband, National Institutes of Health, USA) was used for all morphometric measurements.

### Transmission Electron Microscopy

Larvae collected at early 9 dpf were fixed in 4% glutaraldehyde, 1% paraformaldehyde and 1% sucrose in 0.1 M HEPES at 4°C overnight and embedded in EPON 812 (Serva, Heidelberg, Germany) as per manufacturer’s instructions. Semithin (500 nm) and ultrathin sections (70 nm) were made on an Ultracut UCT microtome (Leica Microsystems, Heidelberg, Germany). Semithin sections were stained with methylene blue. UItrathin sections were placed on copper grids, contrasted with 5% uranyl acetate for 5 min and with Sato’s lead stain for 5 min. Images were acquired with a LIBRA 120 transmission electron microscope (Carl Zeiss GmbH, Oberkochen, Germany) with an anode voltage of 80kV.

### Statistical analysis

GraphPad prism V5.01 (GraphPad Software, CA, USA) was used for all statistical analyses. Gaussian distribution was checked by Kolmogorov-Smirnov testing. If passed, Student’s t-test was used for significance testing and mean was given in the results. For statistical testing of nonparametric data, Mann-Whitney-U-test was applied and median was used. P-values lower 0.05 were considered statistically significant.

## Author Contributions

K.U.I.H., F.S., S.D., M.S. and A.B. established methods and performed experiments. K.U.I.H., F.S., K.E. and N.E. planned experiments, analyzed and interpreted data. W.Z. established the transgenic lines used in the study. All authors reviewed and approved the final version of the manuscript.

The authors declare no conflict of interest.

This article contains supporting information online.

## Acknowledgments

This work was supported by scholarships of the Gerhard Domagk program of the University Medicine Greifswald to K.H., F.S. and S.D. and by a grant of the Federal Ministry of Education and Research (BMBF, grant 01GM1518B, STOP-FSGS) to N.E. The authors thank Mandy Weise and Oliver Zabel for excellent technical assistance.

