## Supplemental Figures for "Prolonged podocyte depletion in larval zebrafish resembles mammalian focal and segmental glomerulosclerosis"

*Address for correspondence:
Prof. Dr. rer. nat. Nicole Endlich
Friedrich-Loeffler Str. 23c, 17487 Greifswald, Germany


### Suppl. Figure 1

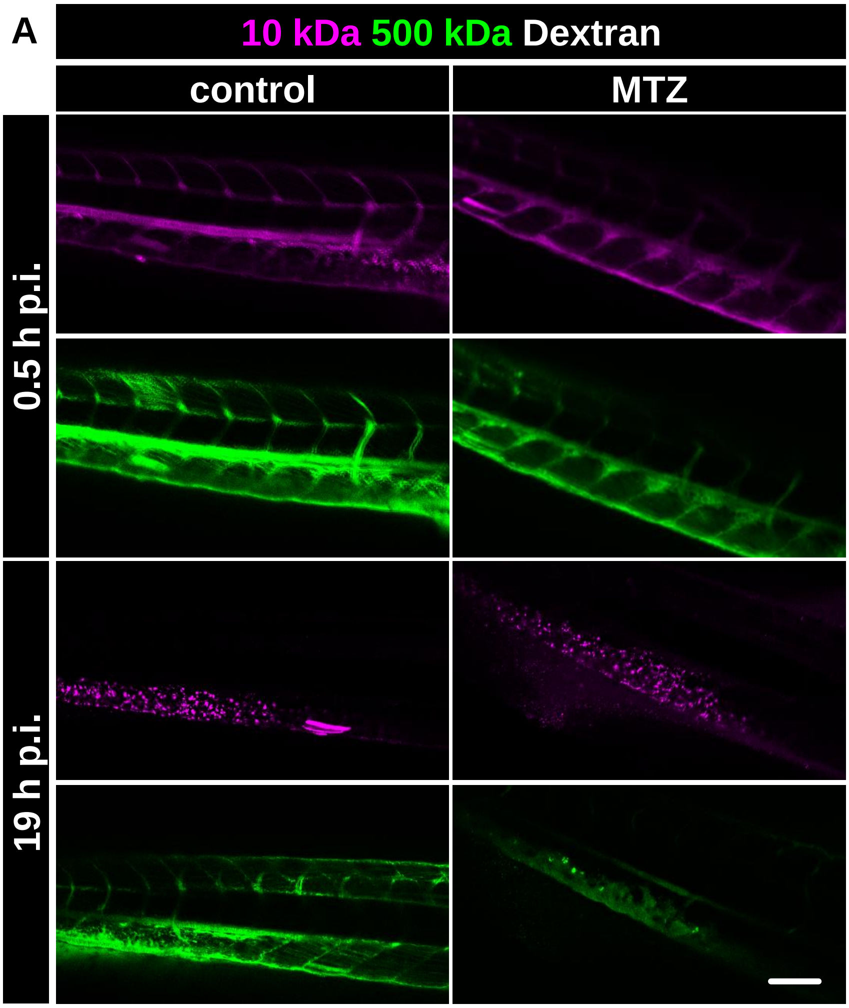


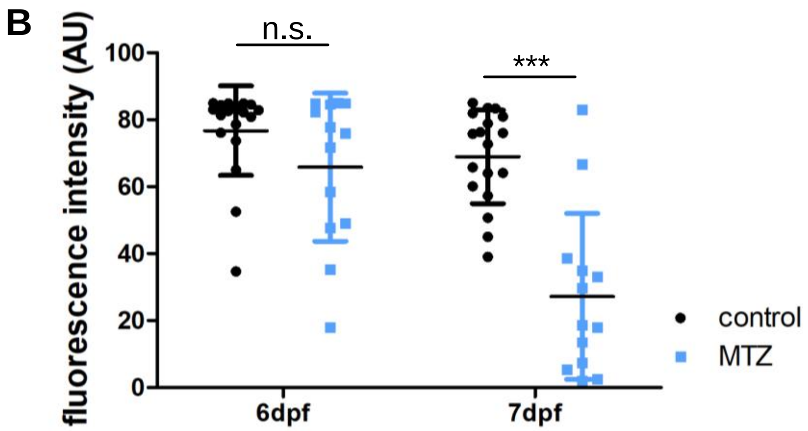


**Supplemental Figure 1**
Panel A shows confocal laser scanning micrographs of dextran-injected larvae 0.5 and 19 h after injection. The scale bar represents 50 µm. 500 kDa dextran is decreased in podocyte-depleted larvae, whereas 10 kDa dextran accumulates in the tissue. As shown in graph B, fluorescence intensity of FITC measured in the caudal vein was statistically significantly lower in podocyte-depleted larvae only 19 h after dextran injection and not directly after injection.

**Suppl. Figure 2**

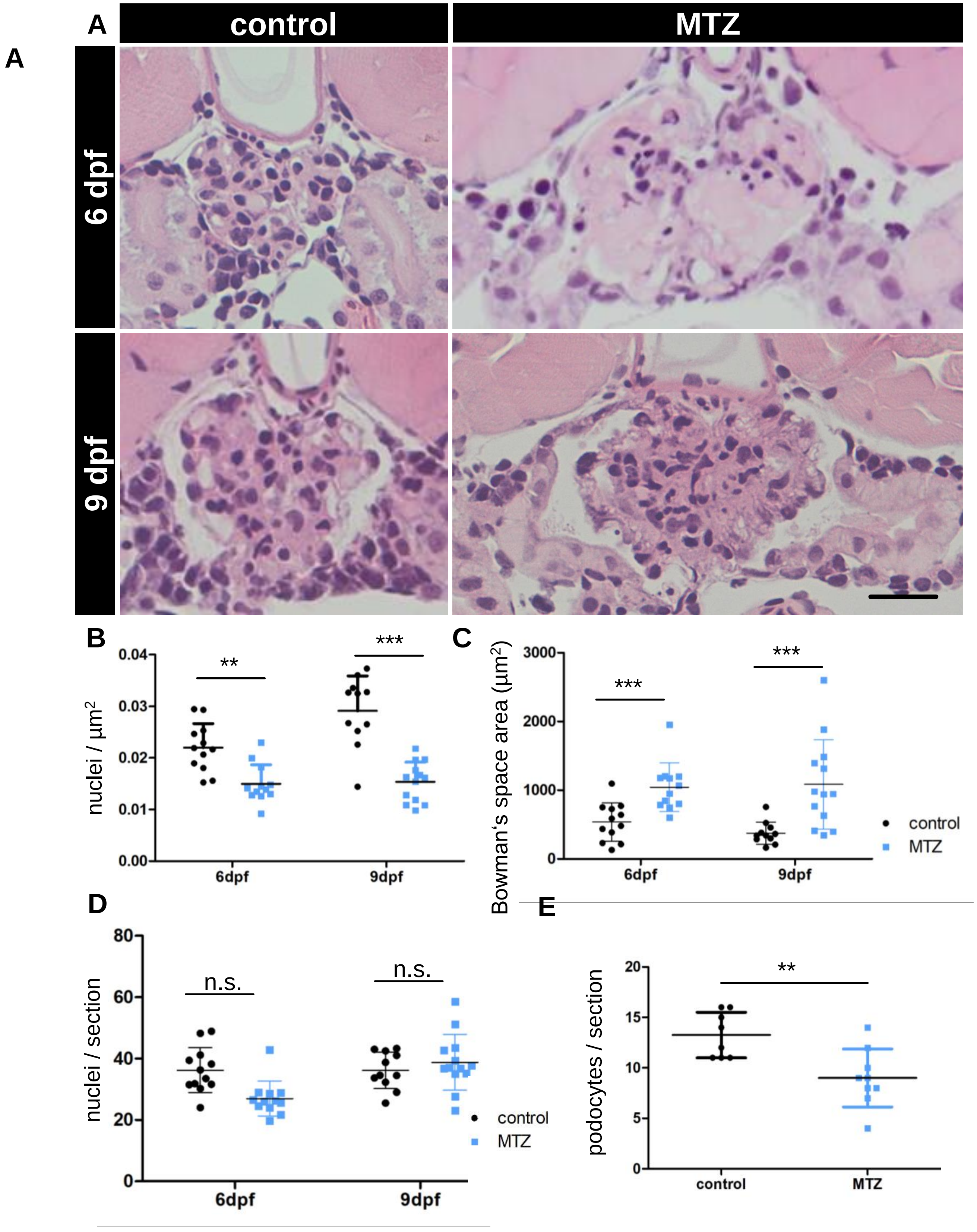

**Supplemental Figure 1**

Panel A shows representative micrographs of H&E-stained plastic sections of podocyte-depleted and control larvae. The scale bar represents 20 µm. As shown in graph B, cell nuclei per µm2 as determined in sections of n=25 podocyte-depleted larvae are significantly reduced compared to n=23 control larvae. Graph C shows a significant increase of Bowman's space area in podocyte-depleted larvae determined by morphometric examination of plastic sections. As shown in graph D, absolute cell counts in cross-sections of the capillary tuft of MTZ-treated larvae show a decrease compared to sections of control larvae at 6 dpf (median 26.11 nuclei per section in MTZ-treated larvae versus 34.92 in controls; p=0.0022; n=48). Until 9 dpf, cell counts recover to a level similar to controls (37.0 nuclei per section in MTZ-treated larvae versus 34.57 in controls; p=0.4173; n=24). Nevertheless, podocyte numbers per glomerular cross section at 9 dpf are still below controls (E).

### Suppl. Figure 3

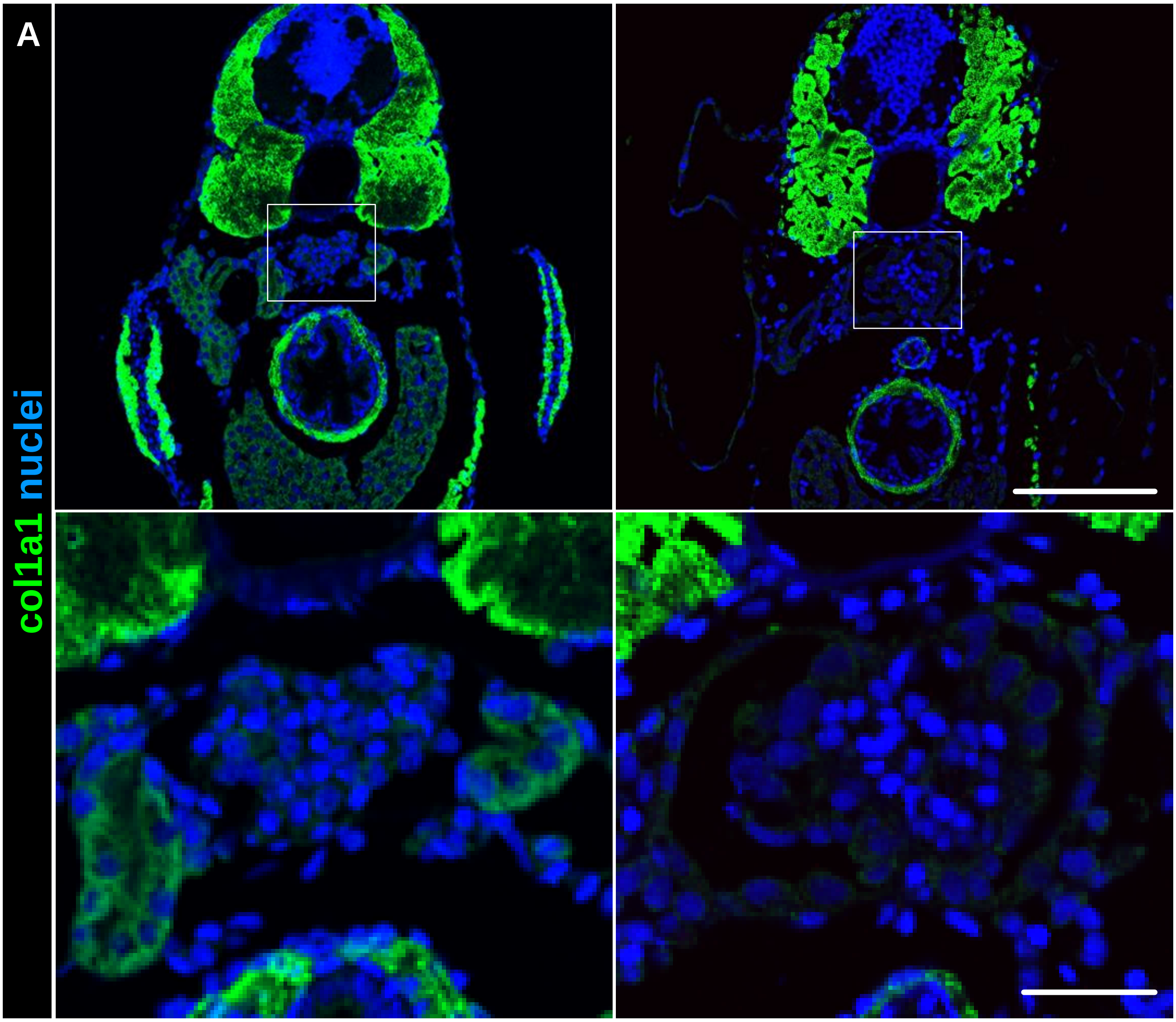


**Supplemental Figure 3**
Representative staining for collagen I alpha chain (col1a1) is shown in panel A with the glomeruli displayed in detail below. No col1a1 positive signal was found in the glomeruli of control or podocyte-depleted larvae. Scale bars represent 100 µm in the upper picture and 25 µm in the lower picture.

### Suppl. Figure 4

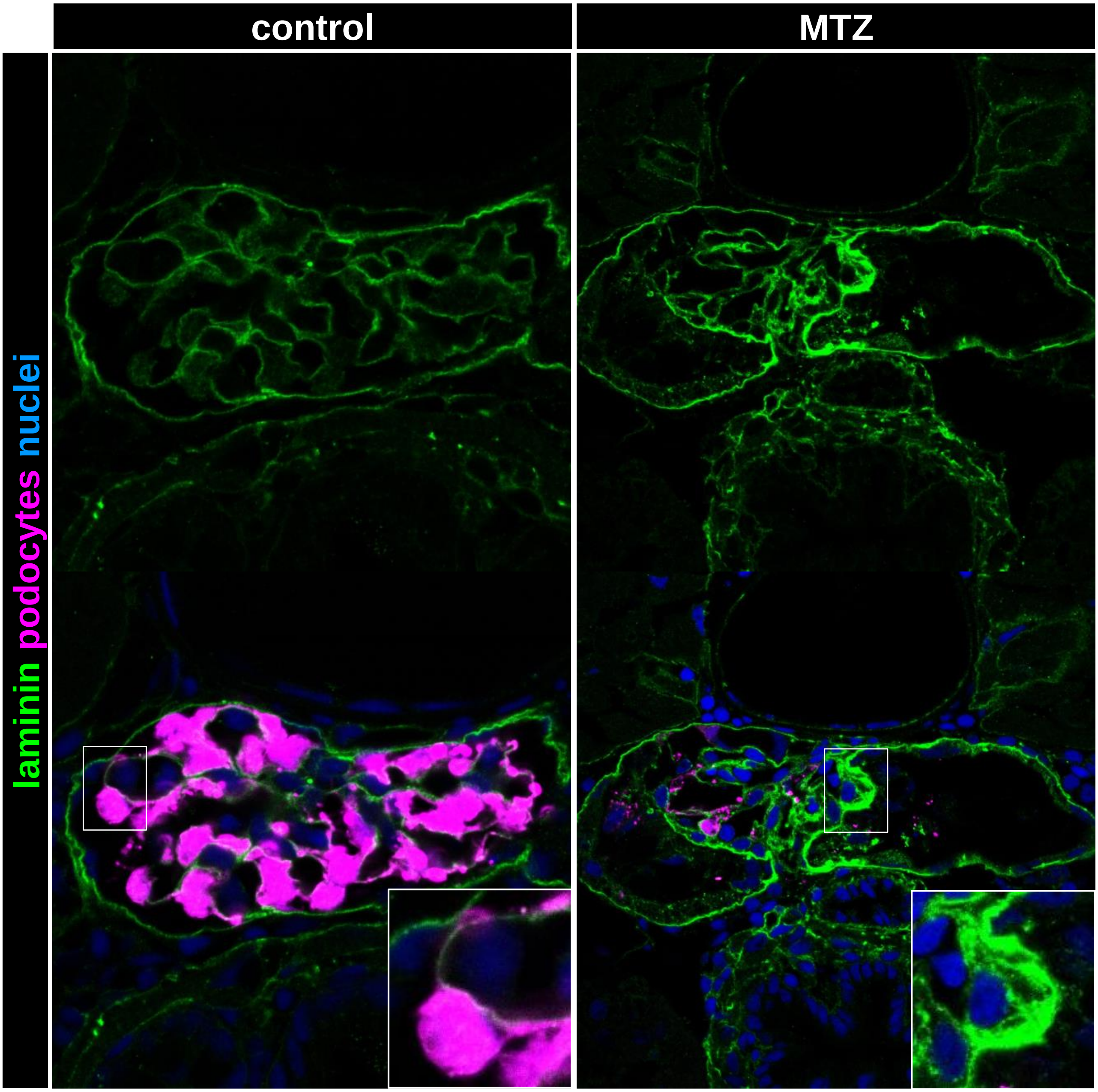


**Supplemental Figure 4**

This figure gives an overview of the laminin staining pattern in 9 dpf larvae. The antibody selectively stains basement membranes as demonstrated on the visceral and parietal basement membrane of the glomerular tuft. Significantly deposition of laminin can be seen after podocyte depletion.

#### Suppl. Figure 5

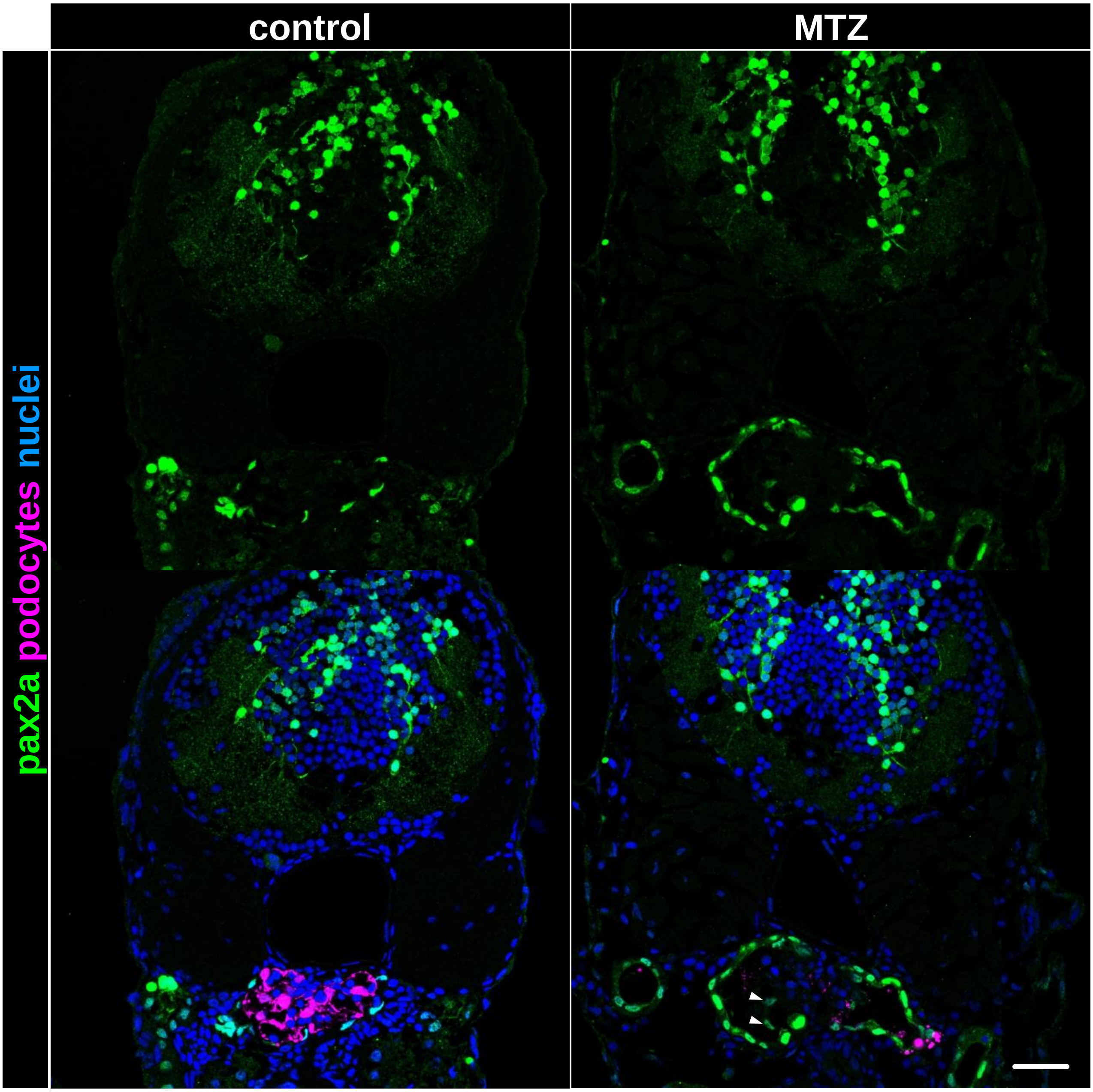


**Supplemental Figure 5**
Immunofluorescence staining for *pax2a* in 9 dpf larvae shows *pax2a* expression in the hindbrain, the neck segment of the proximal tubules, as well as PECs under baseline conditions. After podocyte-depletion, the expression of *pax2a* in PECs is greatly increased. Additionally, large cells on the glomerular tuft express *pax2a* (arrowheads), which could not be found in control specimen. Scale bar represents 25 µm.

Suppl. Figure 6

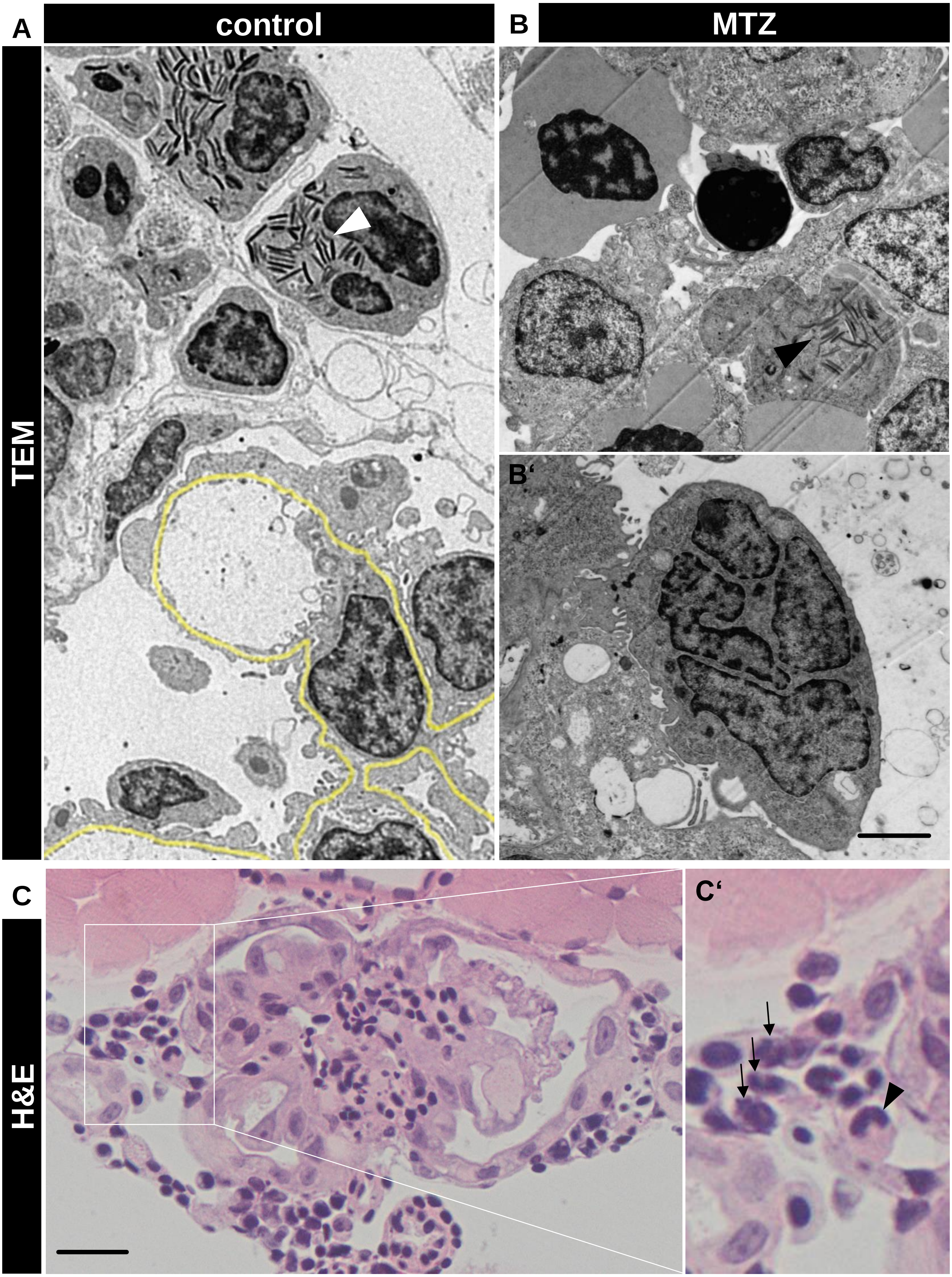

**Supplemental Figure 6**

Transmission electron micrographs of control larvae (A) collected three days after treatment revealed that neutrophils displaying characteristic electron-dense granules (white arrowhead) are located in the interstitium surrounding the glomerulus but not intraglomerular. GBM is highlighted in yellow. Cells with electron-dense granules (arrowheads in B) and a lobate nucleus (B') found in the capillary tuft of podocyte-depleted larvae are shown in detail. Scale bar represents 2 µm. An H&E-stained plastic section of a podocyte-depleted glomerulus is exemplary shown in picture C. Immigrating leukocytes are shown in detail in C'. Black arrows mark neutrophils with the characteristic lobate nucleus. The black arrowhead marks the horseshoe-shaped nucleus of an intracapsular macrophage. Scale bar represents 20 µm.
